# Differential pathogenesis of SARS-CoV-2 variants of concern in human ACE2-expressing mice

**DOI:** 10.1101/2022.04.04.486975

**Authors:** Janhavi Prasad Natekar, Heather Pathak, Shannon Stone, Pratima Kumari, Shaligram Sharma, Komal Arora, Hussin Alwan Rothan, Mukesh Kumar

## Abstract

Severe acute respiratory syndrome coronavirus 2 (SARS-CoV-2) has caused a pandemic resulting in millions of deaths worldwide. Increasingly contagious variants of concern (VoC) have fueled recurring global infection waves. A major question is the relative severity of disease caused by the previous and currently circulating variants of SARS-CoV-2. In this study, we evaluated the pathogenesis of SARS-CoV-2 variants in human ACE-2-expressing (K18-hACE2) mice. Eight-week-old K18-hACE2 mice were inoculated intranasally with a representative virus from the original B.1 lineage, or the emerging B.1.1.7 (alpha), B.1.351 (beta), B.1.617.2 (delta) or B.1.1.529 (omicron) lineages. We also infected a group of mice with the mouse-adapted SARS-CoV-2 (MA10). Our results demonstrate that B.1.1.7, B.1.351 and B.1.617.2 viruses are significantly more lethal than B.1 strain in K18-hACE2 mice. Infection with B.1.1.7, B.1.351 and B.1.617.2 variants resulted in significantly higher virus titers in the lungs and brain of mice compared to the B.1 virus. Interestingly, mice infected with the B.1.1.529 variant exhibited less severe clinical signs and high survival rate. We found that B.1.1.529 replication was significantly lower in the lungs and brain of infected mice in comparison to other VoC. Transcription levels of cytokines and chemokines in the lungs of the B.1.1.529-infected mice were significantly less when compared to those challenged with the B.1.1.7 virus. Together, our data provide insights into the pathogenesis of the previous and circulating SARS-CoV-2 VoC in mice.

## 1. Introduction

SARS-CoV-2 is a positive-sense, single-stranded RNA virus belonging to Betacoronavirus family [1,2]. Since the emergence of SARS-CoV-2 in late 2019, several new variants of concern (VoC) alpha (B.1.1.7 lineage), beta (B.1.351 lineage), gamma (P.1 lineage), delta (B.1.617.2 lineage) and omicron (B.1.1.529 lineage) have fueled recurring global infection waves. These variants have been termed VoC because of the higher risk due to their possible enhanced transmissibility, disease severity, immune escape, and increased adaptation to new hosts [3–8]. Mutations occurring in the spike protein are of major concern due to the role of this glycoprotein in mediating virus entry and as the major target of neutralizing antibodies [3,9–11]. The lineage B.1.1.7 was first identified in the United Kingdom, lineage B.1.351 was discovered in South Africa, and lineage B.1.617.2 was described in India. Most recently, omicron (B.1.1.529) VoC that emerged in South Africa is estimated to have been responsible for majority of infections worldwide. The B.1.1.7 variant has mutations in the receptor binding domain (RBD) region, including N501Y, 69/70 deletion, and P681H near the S1/S2 furin cleavage site [7,12–14]. The B.1.351 variant has eight mutations, of which the three most notable mutations are K417N, E484K, and N501Y in the spike protein [3,7,9,15]. The B.1.617.2 variant has three unique mutations, E156del/R158G in the N-terminal domain and T478K in RBD of the spike protein. The B.1.1.529 variant has an unusually large number of mutations in the spike protein, including 30 amino acid substitutions, three short deletions, and one insertion [16–18].

The main goal of this study was to compare the SARS-CoV-2 variants replication and pathogenesis in K18-hACE2 mice. K18-hACE2 is a transgenic mouse model which expresses human ACE2 driven by the human cytokeratin 18 promoter. K18-hACE2 mice is a well-established model for SARS-CoV-2 studies that supports virus replication in the respiratory and central nervous systems, resulting in elevated chemokine and cytokine levels and significant tissue pathologies [19–21]. Our results demonstrate that B.1.1.7 (alpha), B.1.351 (beta) and B.1.617.2 (delta) variants are more virulent than the original SARS-CoV-2 B.1 Wuhan strain in K18-hACE2 mice. Infection with B.1.1.7, B.1.351 and B.1.617.2 variants resulted in significantly high virus titers in the lungs and brain of mice compared to the B.1 virus. Interestingly, the replication capacity of the omicron variant was significantly lower than other VoC. Mice infected with the B.1.1.529 virus exhibited high survival rate and had a lower virus load in the lungs and brain compared to mice infected with B.1.1.7, B.1.351 and B.1.617.2 viruses. In addition, transcription levels of cytokines and chemokines in the lungs of the B.1.1.529-infected mice were significantly lower when compared to those challenged with the B.1.1.7 virus. Together, our data provide insights into the pathogenesis of the previous and circulating SARS-CoV-2 VoC in mice.

## 2. Materials and Methods

### 2.1. In vivo mouse challenge experiments

*In vivo* mouse experiments involving infectious SARS-CoV-2 were performed in Animal Biosafety Level 3 laboratory and strictly followed the approved standard operation procedures. The protocol was approved by the Georgia State University Institutional Animal Care and Use Committee (Protocol number A20044). Hemizygous K18-hACE2 mice (2B6.Cg-Tg (K18-ACE2)2Prlmn/J) were obtained from The Jackson Laboratory. Eight-week-old hemizygous K18-ACE2 mice were inoculated intranasally with PBS (mock) or 10^4^ plaque-forming units (PFU) of SARS-CoV-2 as described previously [8,21,22]. We used B.1 Wuhan virus (BEI# NR-52281), B.1.1.7 virus (BEI# NR-54000), B.1.351 virus (BEI# NR-54008), B.1.617.2 virus (Northwestern Reference laboratory, Clinical isolate #2333067), B.1.1.529 virus (BEI# NR-56461) and MA10 virus (BEI# NR-55329). Roughly equal numbers of male and female mice were used. Animals were weighed and their appetite, activity, breathing, and neurological signs were assessed twice daily. In independent experiments, mice were inoculated with PBS (Mock) or SARS-CoV-2 intranasally, and on days 3 and 5-7 after infection, animals were anesthetized using isoflurane and perfused with cold PBS. The lungs and brains were collected and flash-frozen in 2-methylbutane (Sigma, St. Louis, MO, USA) for further analysis, as described below [8,23].

### 2.2. Infectious virus titration by plaque assays

Tissues harvested from virus-inoculated animals were weighed and homogenized in a bullet blender (Next Advance, Averill Park, NY, USA) using stainless steel or zirconium beads, followed by centrifugation and titration. Virus titers in tissue homogenates were measured by plaque assay using Vero E6 cells [8]. To titrate the infectious virus, tissue homogenates were 10-fold serially diluted with DMEM and applied to monolayered Vero E6 cells for 1 hour. After inoculation, cells were washed once before overlaid with 1% low-melting agarose. Cells were further incubated for 48 hours and stained with neutral red for visualizing plaque formation [8,24].

### 2.3. RNA extraction and quantitative RT-PCR

Total RNA was extracted from the lungs using a Qiagen RNeasy Mini kit (Qiagen, Germantown, MD, USA). The cDNA was synthesized from RNA using an iScript^™^ cDNA Synthesis Kit (Bio-Rad). The qRT-PCR was used to determine the expression levels of IL-6 and CCL-2 as described previously [22,24]. The fold-change in infected samples compared to control samples was calculated after normalizing to the housekeeping GAPDH gene [22,23]. The primer sequences used for qRT-PCR are listed in Table 1.

**Table 1.**
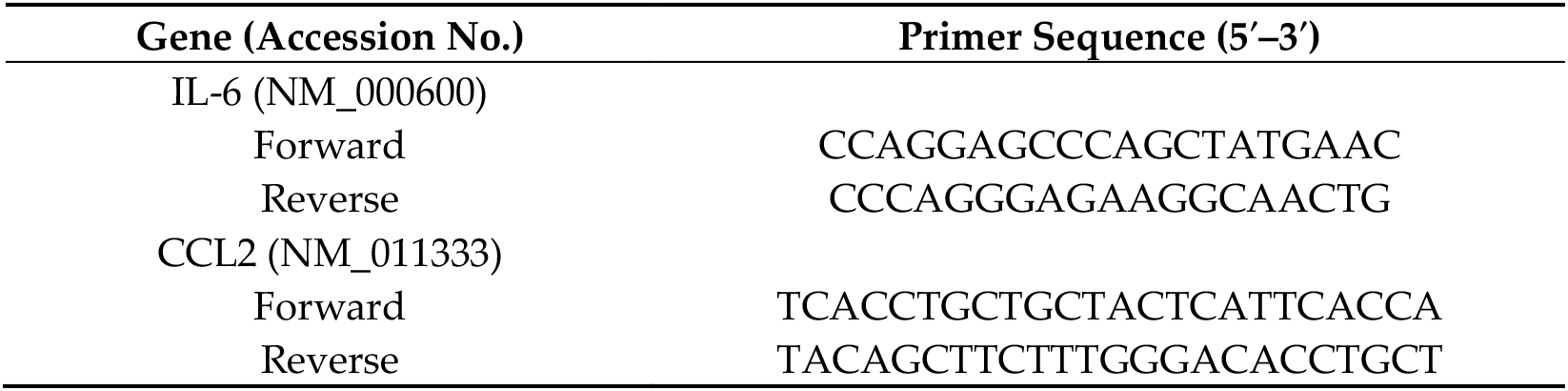
Primer sequences used for qRT-PCR.

### 2.4. Statistical Analysis

GraphPad Prism 8.0 was used to perform a Kaplan Meier log-rank test to compare survival curves. For body weight changes, two-way analysis of variance (ANOVA) with the post hoc Bonferroni test was used to calculate values of p. Mann–Whitney tests and unpaired student *t*-tests were used to calculate the *p* values of the difference between viral titers and immune responses, respectively. Differences of *p* < 0.05 were considered significant.

## 3. Results

### 3.1. Clinical disease progression of K18-hACE2 mice infected with SARS-CoV-2 VoC

To evaluate the pathogenicity of the original B.1 lineage and emerging SARS-CoV-2 lineages, K18-hACE2 mice were infected intranasally with a representative virus from the original B.1 lineage, or the emerging B.1.1.7 (alpha), B.1.351 (beta), B.1.617.2 (delta) and B.1.1.529 (omicron) lineages. We also infected a group of mice with the mouse-adapted SARS-CoV-2 (MA10) [25]. Animals were monitored for clinical signs and survival. The mock-infected mice remained healthy throughout the study period. While the infectious dose of 10^4^ plaque-forming units (PFU) of B.1 virus resulted in 75% mortality, mortality in B.1.1.7-, B.1.351-, B.1.617.2-infected mice was 100% (Figure 1A). The median survival time of mice infected with alpha, beta and delta variants was also shorter than in the B.1-infected mice. Statistically, mouse survival for B.1 virus was significantly higher than alpha, beta and delta variants. K18-hACE2 mice infected with the MA10 virus showed a faster disease progression and severity after infection compared to all SARS-CoV-2 clinical isolates. Interestingly, infection with the B.1.1.529 (omicron) virus resulted in only 50% mortality with extended survival time in mice (Figure 1A). There was a significant difference between the survival of the B.1.1.529 challenged mice compared to the other VoC at the same dose.

**Figure 1.**
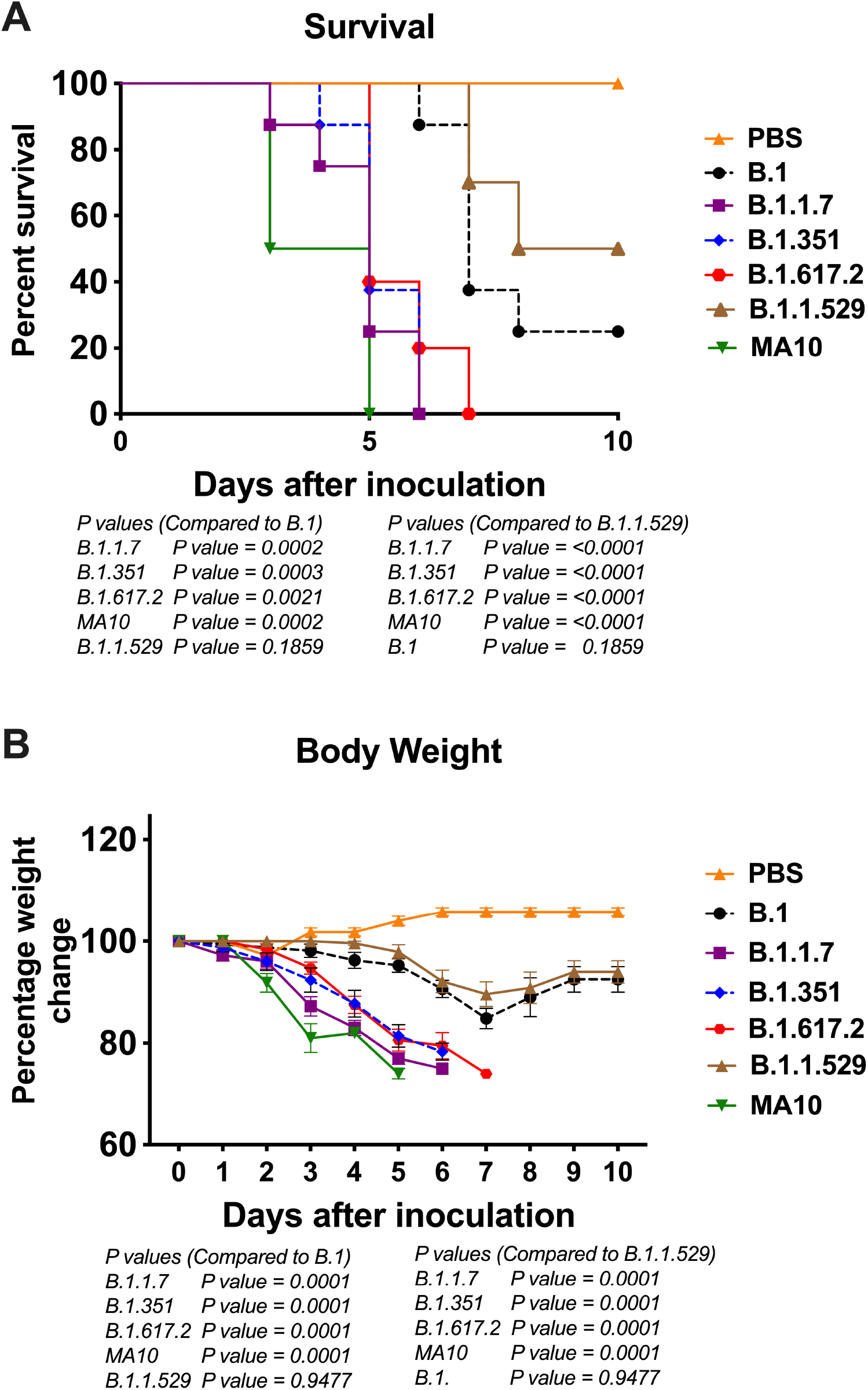
Analysis of Survival and body weight in K18-hACE2 mice following infection with SARS-CoV-2 VoC. Eight-week-old K18-hACE2 mice were inoculated intranasally with PBS (mock) or 10^4^ PFU of B.1 and individual variants (n = 10-15 mice per group). **(A)** Survival curve **(B)** Body weight change in percentage. Values are the mean ± SEM.

As early as 3 days after infection, mice inoculated with B.1.1.7 and B.1.351 virus began to lose body weight and showed signs of infection. By 6 days, all mice infected with B.1.1.7 or B.1.351 virus died after losing 20% body weight and experiencing severe symptoms (Figure 1B). Mice infected with the B.1.617.2 also lost significant body weight, and all succumbed to death by day 7 after infection. Statistically, body weight loss for B.1.1.7, B.1.351 and B.1.617.2 viruses was significantly higher than the B.1 virus. Compared to the other VoC, the body weight loss of mice infected by the B.1.1.529 virus was significantly milder with onset time at a later stage during the infection (Figure 1B).

### 3.2. Viral load in K18-hACE2 mice infected with SARS-CoV-2 VoC

To evaluate virus replication in the tissues, groups of 3-7 mice were euthanized at 3- and 5-7-days after infection, and the lungs and brain were collected. Viral infectivity titers in the tissues were measured by plaque assay. A median infectious virus titer of 5 × 10^5^ PFU/g was detected at day 3 after infection in the lungs from the animals infected with the B.1 virus. Compared to the B.1 virus, infection with B.1.1.7, B.1.351 and B.1.617.2 viruses resulted in significantly higher levels of infectious virus in the lungs at 3 days after infection (Figure 2A). On 5-7 days after infection, mice infected with B.1.1.7, B.1.351 and B.1.617.2 viruses sustained significantly high levels of viral load in the lungs compared to the B.1 virus (Figure 2B). In contrast, the replication of the B.1.1.529 virus was dramatically reduced in comparison with that of B.1.1.7, B.1.351 and B.1.617.2 viruses, despite using the same inoculation titers for virus challenge. The levels of infectious virus in the lungs of B.1.1.529-infected mice were approximately 100-fold lower than in the animals infected with other VoC at both 3 and 5-7 days after infection (Figures 2A and 2B).

**Figure 2.**
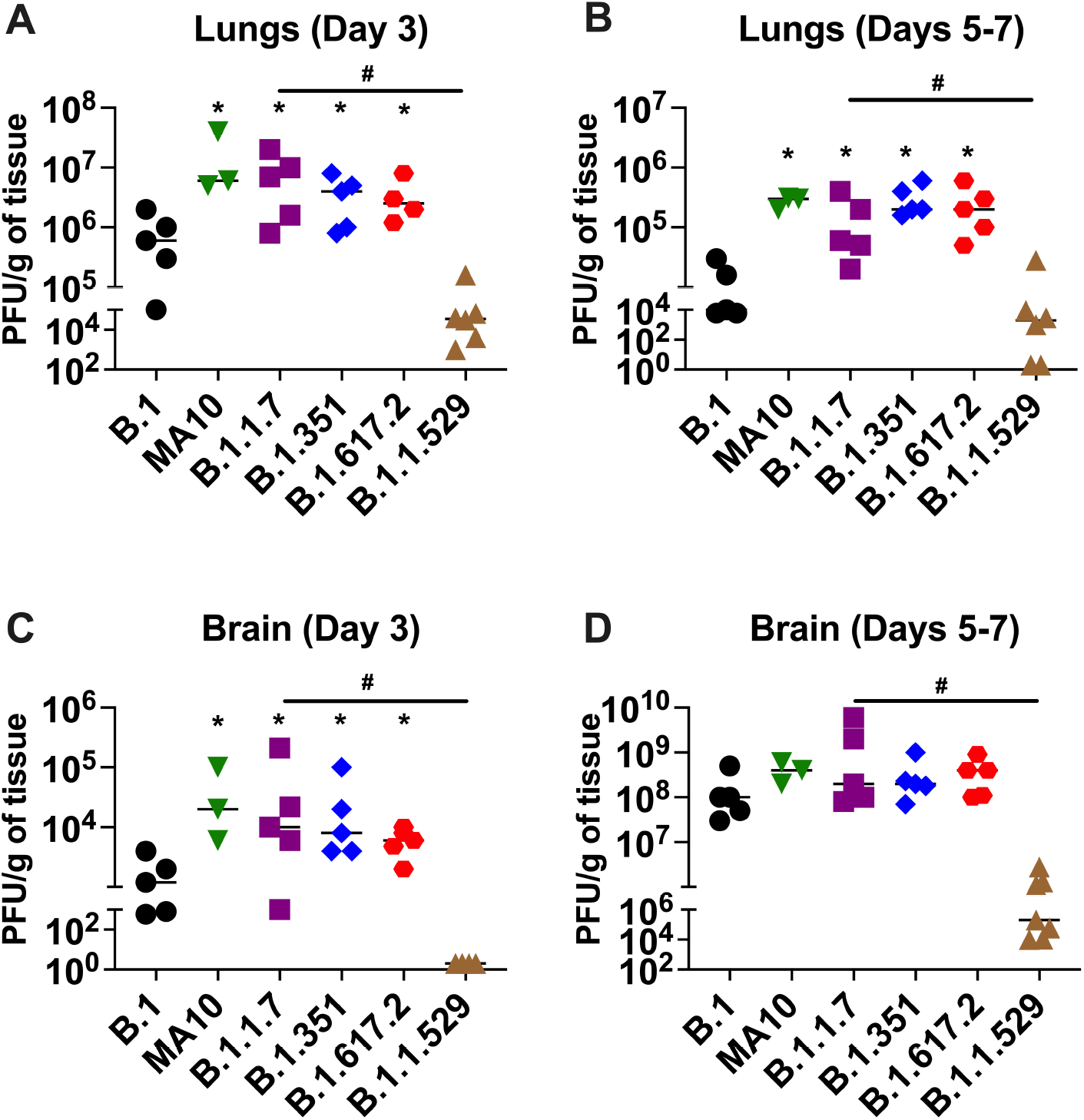
Replication of SARS-CoV-2 VoC in the lungs and brain. Eight-week-old K18-hACE2 mice were inoculated intranasally with 10^4^ PFU of SARS-CoV-2 or variants. Groups of 3-7 mice were euthanized on day 3 and days 5-7 after infection and tissues were collected. Virus titers were analyzed in the lungs and brain by plaque assay. The data are expressed as PFU/g of tissue. **(A)** virus titer in day 3 lungs, **(B)** virus titer in days 5-7 lungs, **(C)** virus titer in day 3 brain, and **(D)** virus titer in days 5-7 brain. Each data point represents an individual mouse. *, *p* < 0.05 (compared to B.1 virus); #, *p* < 0.05 (compared to B.1.1.529 virus).

In the brain, mice infected with the B.1 virus exhibited significantly lower levels of infectious virus than B.1.1.7-, B.1.351- and B.1.617.2-infected animals at day 3 after infection (Figure 2C). At 5-7 days after infection, the viral loads were similar in animals infected with B.1, B.1.1.7, B.1.351 and B.1.617.2 viruses (Figure 2D). Consistent with the lung data, infectious virus in the brains harvested from the B.1.1.529-infected mice was also significantly reduced compared to that of the other groups. We did not detect any infectious virus in the B.1.1.529-infected mice at day 3 after infection. On 5-7 days after infection, the viral load was approximately 1,000-fold higher in the B.1.1.7-, B.1.351- and B.1.617.2-infected mice than the B.1.1.529-infected mice (Figures 2C and 2D). A relative decrease in viral replication correlate with the observed decline in clinical and pathological severity and recovery seen in the omicron virus-infected mice.

### 3.2. Inflammation in the lungs following infection with SARS-CoV-2 VoC

The excessive inflammatory host response to SARS-CoV-2 infection contributes to pulmonary pathology and the development of respiratory distress in some COVID-19 patients [26,27]. We next quantified the gene expression of IL-6 and CCL-2 in the lungs of K18-hACE2 mice at day 3 after infection. Gene expression changes in the lungs of infected mice, compared with the mock-infected controls, were analyzed by qRT-PCR. Mice inoculated with the omicron variant had lower mRNA levels of IL-6 and CCL-2 compared to those inoculated with the alpha variant. As shown in figure 3, B.1.1.7 infection resulted in an approximately 100-fold increase in IL-6 and CCL-2 mRNA expression. In contrast, IL-6 and CCL-2 mRNA levels increased by only 10–20-fold in the lungs of the B.1.1.529-infected mice, suggesting attenuated inflammation (Figure 3).

**Figure 3.**
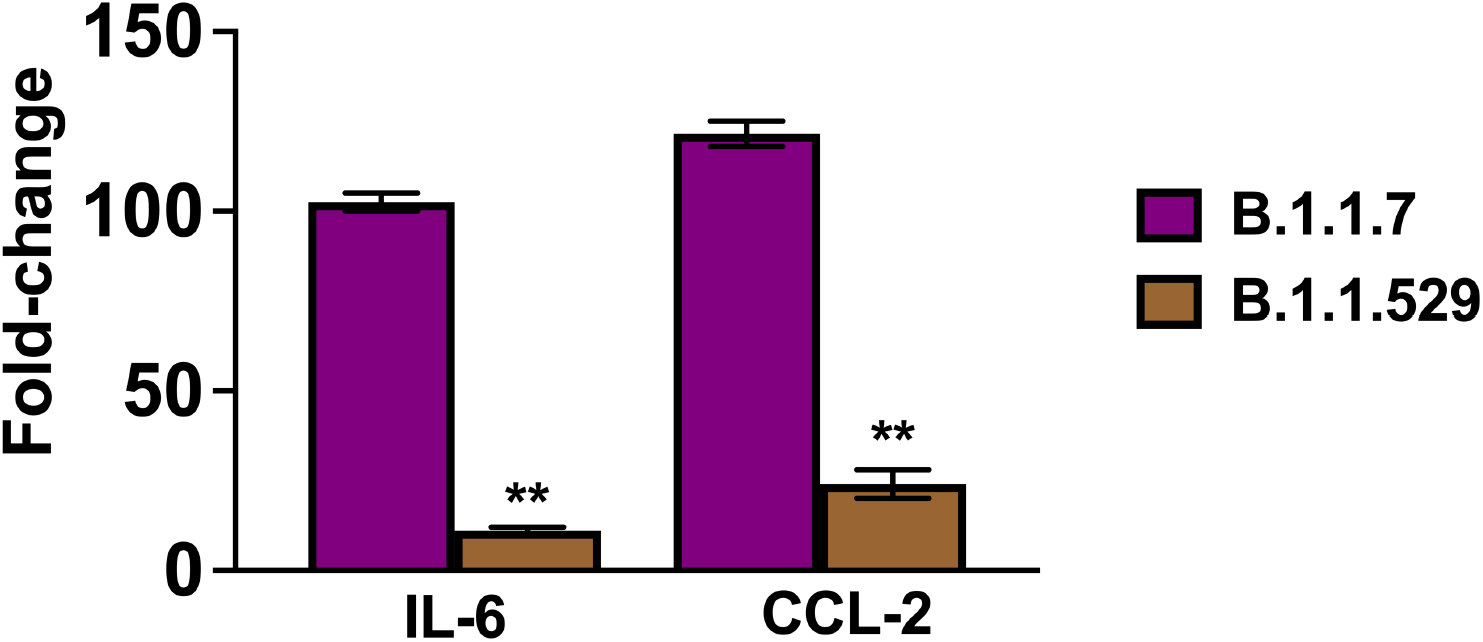
Analysis of inflammatory response in the lungs. Eight-week-old K18-hACE2 mice were inoculated intranasally with PBS (mock) or 10^4^ PFU of B.1.1.7 and B.1.1.529 viruses (n = 5-7 mice per group). Lungs and brain were harvested after extensive perfusion with PBS and RNA was extracted. The mRNA levels of IL-6 and CCL-2 were determined by qRT-PCR. Values are the mean ± SEM. **, *p* < 0.001

## 4. Discussion

To date, most K18-hACE2 mouse studies have utilized original viral strains of SARS-CoV-2 and few studies have been performed with emergent VoC. In this study, we investigated the pathogenicity of SARS-CoV-2 variants in K18-hACE2 mice. Our results demonstrate that the pathogenicity of SARS-CoV-2 in K18-hACE2 mice is VoC-dependent and highest for alpha, beta and delta. We found significantly high virus titers in the lungs and brain of mice infected with B.1.1.7, B.1.351 and B.1.617.2 variants compared to the B.1 lineage. In contrast, the omicron variant replicated significantly less efficiently than other SARS-CoV-2 variants in mice. In comparison with alpha, beta, and delta variant, omicron variant results in the less body weight loss and mortality rate. This is also reflected by less extensive inflammatory processes in the lungs of the B.1.1.529-infected mice.

SARS-CoV-2 evolves rapidly with accumulation of mutations in viral genome, giving rise to multiple variants of concern (VoC) [4,5,10]. Among all mutations, N501Y is the most critical because it involves amino acid residues that account for the tight binding of RBD of the SARS-CoV-2 and ACE2 receptors on the host cell surface [7,10,11]. B.1.1.7 and B.1.351 both lineages contain N501Y in RBD in addition to widespread D614G in spike protein, while the B.1.351 also has two additional mutations; K417N and E484K [10,12,13,15]. Both N501Y and D614G have been shown to increase RBD binding to ACE2 and promote virus entry and replication in humans and animal models [3,7–11]. These mutations in the RBD of the spike protein may have enhanced the binding affinity for the ACE2 receptor, thereby allowing the variants to replicate more efficiently in mice. Indeed, our results demonstrate that SARS-CoV-2 variants B.1.1.7 and B.1.351 containing N501Y and E484K mutations display a substantially severe pathogenicity in K18-hACE2 mice. B.1.1.7 and B.1.351 caused lethal disease in K18-hACE2 mice accompanied with high tittered viral burdens in the lungs and brain. Our results corroborated with the recent findings that alpha and beta variants were able to cause enhanced disease in K18-hACE2 mice [6,7,14,28]. The B.1.617.2 (delta) lineage does not harbor the N501Y substitution in the spike protein but does have additional mutations within the spike protein which diverge from other VoC. In the B.1.617.2 lineage, two modifications, namely L452R and T478K in the RBD, increase the interaction with ACE2 with the highest binding affinity [29,30], which may contribute to increase in pathogenicity observed in mice.

Despite carrying the highest number of mutations that might allow more efficient binding to ACE2, we observed mild disease in mice infected with the omicron variant as compared with other SARS-CoV-2 variants. Our results agree with recent studies that demonstrated attenuated replication of the omicron variant in mice and hamsters [16,18]. Recently, *in vitro* studies have shown that the omicron variant is inefficient in transmembrane serine protease 2 (TMPRSS2) usage in comparison to that of previous variants [31,32]. It is possible that attenuated replication of the omicron variant in human cells and mice is due to a reduced efficiency in host proteases cleavage by changes in the spike protein. Epidemiological data also suggest that the omicron virus causes a less severe pathology in humans than the ancestral strains and the other VoC [29]. Overall, our study demonstrates that SARS-CoV-2 pathogenicity in K18-hACE2 mice is VoC-dependent and highest for alpha and beta, and lowest for the omicron variant. These results will be valuable for understanding the pathogenesis of emerging SARS-CoV-2 variants.

## Author Contributions

Conceptualization, M.K.; methodology, J.P.N., H.P.; S.S., P.K., H.A.R., K.A., M.K.; validation, J.P.N., H.P.; S.S., P.K., S.S., K.A., M.K.; formal analysis, P.N., H.P.; S.S., P.K., H.A.R., S.S., K.A., M.K.; resources, M.K.; writing–original draft preparation, J.P.N., H.P.; K.A., M.K.; writing–review and editing, K.A., M.K.; funding acquisition, M.K. All authors have read and agreed to the published version of the manuscript.

## Institutional Review Board Statement

The animal study protocol was approved by the Georgia State University Institutional Animal Care and Use Committee (Protocol number A20044).

## Informed Consent Statement

Not applicable.

## Data Availability Statement

Not applicable.

## Acknowledgments

We thank members of the GSU High Containment Core and the Department for Animal Research for assistance with the experiments.

